# Genetic models suggest single and multiple origins of dihydrofolate reductase mutations in *Plasmodium vivax*

**DOI:** 10.1101/2020.09.18.303586

**Authors:** Ayaz Shaukat, Qasim Ali, Lucy Raud, Abdul Wahab, Taj Ali Khan, Imran Rashid, Muhammad Rashid, Mubashir Hussain, Mushtaq A. Saleem, Neil D. Sargison, Umer Chaudhry

## Abstract

Pyrimethamine was first introduced for the treatment of malaria in Asia and Africa during the early 1980s, replacing chloroquine, and has become the first line of drugs in many countries. In recent years, development of pyrimethamine resistance in *Plasmodium vivax* has become a barrier to effective malaria control strategies. Here, we describe the use of meta-barcoded deep amplicon sequencing technology to assess the evolutionary origin of pyrimethamine resistance by analysing the flanking region of dihydrofolate reductase (*dhfr*) locus. The genetic modelling suggests that 58R and 173L single mutants and 58R/117N double mutants are present on a single lineage; suggesting a single origin of these mutations. The triple mutants (57L/58R/117N, 58R/61M/117N and 58R/117N/173L) share the lineage of 58R/117N, suggesting a common origin. In contrast, the 117N mutant is present on two separate lineages suggesting that there are multiple origins of this mutation. We characterised the allele frequency of the *P. vivax dhfr* locus. Our results support the view that the single mutation of 117N and double mutations of 58R/117N arise commonly, whereas the single mutation of 173L and triple mutations of 57L/58R/117N, 58R/61M/117N and 58R/117N/173L are less common. Our work will help to inform mitigation strategies for pyrimethamine resistance in *P. vivax*.

## 1. Introduction

The estimated 228 million cases of malaria and 405,000 deaths in 2018 represent a huge global public health burden (Poostchi et al., 2018). Sustainable malaria prevention requires reliable surveillance of drug resistance (Ding et al., 2013). With increasing reports of the pyrimethamine resistance-associated mutations in *Plasmodium vivax*, it is important to consider effective prevention and control strategies; while reducing the risk of further development of resistance mutations (Petersen et al., 2011; Shaukat et al., 2019). Pyrimethamine inhibits dihydrofolate reductase (*dhfr*) enzymes of *P. vivax*, thereby blocking pyrimidine biosynthesis, leading to interruption of DNA synthesis (Eldin de Pécoulas et al., 1998). Single nucleotide polymorphisms (SNPs) in the *P. vivax dhfr* locus have been associated with pyrimethamine resistance. Resistance-associated SNPs have been found at codons F57L/I (TT<u>C</u>-TT<u>A</u>/<u>A</u>T<u>A</u>), S58R (AG<u>C</u>-AG<u>A</u>), T61M (A<u>C</u>G-A<u>T</u>G), S117N/T (A<u>G</u>C-A<u>A</u>C/A<u>C</u>C) and I173L/F (A<u>T</u>T-<u>C</u>TT/<u>T</u>TT) (de Pecoulas et al., 1998; Lee et al., 2010).

Detailed knowledge of pyrimethamine resistance has been demonstrated by highlighting the origins of *Plasmodium falciparum dhfr* resistance mutations in the endemic regions (Cortese et al., 2002; Lumb et al., 2009; McCollum et al., 2008; Nair et al., 2003; Nash et al., 2005). There are few studies examining the evolutionary origins of the mutations in *dhfr* locus causing pyrimethamine resistance in *P. vivax* (Alam et al., 2007; Hawkins et al., 2008a). Therefore a better understanding of the evolution of pyrimethamine resistance mutations in *P. vivax* is needed to inform sustainable control of malaria in endemic regions (Shaukat et al., 2019).

We have develop metabarcoded deep amplicon sequencing platforms to investigate the evolutionary origins of drug resistance mutations in various human and livestock parasites (Ali et al., 2019; Sargison et al., 2019; Shaukat et al., 2019). Here, we describe the application of this method to *P. vivax* isolates from Pakistan. Our aims were to explore the evolutionary origins of *dhfr* locus SNPs conferring pyrimethamine resistance and to investigate the allele frequencies of pyrimethamine resistance-conferring mutations.

## 2. Materials and Methods

### 2.1. Parasite material, genomic DNA preparation and species identification

Blood samples were collected from symptomatic patients seeking malaria diagnosis, who had been referred to the Chughtai diagnostic laboratory in the Punjab province of Pakistan. The study was approved by the Institutional Review Board of the University of Central Punjab, Pakistan (UCP-30818). The samples were taken during the ‘peak malaria season’ (April to October) between 2017 and 2019 by trained paramedical workers under the supervision of local collaborators. The study included patients of all age groups with malaria symptoms including vomiting, fever, headache, chills, sweats, nausea and fatigue. The blood samples were collected following the written consent of the patients. 4% Giemsa-stained smears were routinely made from each sample for examination under oil immersion for the diagnosis of malaria. 50 μL aliquots of blood from malaria positive patients were used for gDNA extraction according to the protocols described in the TIANamp blood DNA kit (Beijing Biopeony Co. Ltd) and sent to the Roslin Institue for PCR amplification, Illumina Miseq run and bioinformatics analysis. A ‘haemoprotobiome’ high-throughput sequencing tool was used to confirm the presence of *Plasmodium* species described by Wahab et al. (2020). In this study, malaria positive field samples were examined to identify the species of *Plasmodium* involved in the infection. Overall, the prevalence of *P. vivax* was 69.8%, *P. falciparum* 29.5% and mixed infection 0.7% (Wahab et al., 2020).

### 2.2. PCR amplification and Illumina Mi-Seq run

A 468 bp fragment of the *P. vivax dhfr* locus was amplified from the samples identified by Wahab et al. (2020). The primer sets, adapter/barcoded PCR amplification conditions and magnetic bead purification were previously described by Shaukat et al. (2019). 10 μl of each barcoded PCR product was combined to make a pooled library. At least six pooled libraries were run on agarose gel electrophoresis to separate PCR products. *dhfr* products were excised from the gel using commercial kits (QIAquick Gel Extraction Kit, Qiagen, Germany). 20 μl of eluted DNA was then purified using AMPure XP Magnetic Beads (1X) (Beckman Coulter, Inc.), before being combined into a single purified DNA pool library. The library was measured with KAPA qPCR library quantification kit (KAPA Biosystems, USA) and then run on an Illumina MiSeq Sequencer using a 600-cycle pair-end reagent kit (MiSeq Reagent Kits v2, MS-103-2003) at a concentration of 15nM with the addition of 15% Phix Control v3 (Illumina, FC-11-2003) previously described by Shaukat et al. (2019).

### 2.3. Bioinformatics data handling

The post-run processing separates all the sequences by sample via the recognised barcoded indices and generates FASTQ files. The MiSeq data analysis was performed with a bespoke pipeline using Mothur v1.39.5 software (Kozich et al., 2013; Schloss et al., 2009) with modifications in the standard operating procedures of Illumina Mi-Seq in the Command Prompt pipeline previously described (Shaukat et al., 2019). Briefly, raw paired read-ends were run in the ‘make.contigs’ command to combine the two sets of reads from each sample. The command extracts sequences and quality score data from the FASTQ files, creating the complement of the forward and reverse reads and combined them into contigs. After removing long, or ambiguous sequence reads (>468 bp) using the ‘screen.seqs’ command, the data were aligned with the *P. vivax dhfr* reference sequence library using the ‘align.seqs’ command. Sequences that did not match with the *P. vivax dhfr* reference sequence library were removed and the ‘summary.seqs’ command. The sequence reads were further run on the ‘screen.seqs’ command to generate the *P. vivax dhfr* FASTQ file. Once the sequence reads were classified as *P. vivax*, a count list of the consensus sequences of each sample was created using the ‘unique.seqs’ command. The count list was further used to create FASTQ files of the consensus sequences of each sample using the ‘split.abund’ command to sort data into groups of rare and abundant based on the cut-off value, followed by the ‘split.groups’ command. Those samples yielding more than 1000 reads (implying sufficient gDNA for accurate amplification) were included in the cut-off value.

### 2.4. Bioinformatic analysis

*P. vivax dhfr* sequences generated from the count list of the consensus sequences were edited and aligned in Geneious Prime 2020.1 software (Kearse et al., 2012). These consensus sequences were used for the calculation of the relative allele frequencies of *dhfr* resistance-associated mutations. To achieve this, *P. vivax dhfr* isolates generated from the count list of the consensus sequences were first assigned to relevant susceptible or resistance mutations based on known SNPs at codons F57L/I (TT<u>C</u>-TT<u>A</u>/<u>A</u>T<u>A</u>), S58R (AG<u>C</u>-AG<u>A</u>), T61M (A<u>C</u>G-A<u>T</u>G), S117N/T (A<u>G</u>C-A<u>A</u>C/A<u>C</u>C) and I173L/F (A<u>T</u>T-<u>C</u>TT/<u>T</u>TT). Allele frequencies were calculated by dividing the number of sequences reads of each isolate that contained these mutations by the total number of reads (R Core Team, 2013; package ggplot2).

For the genetic diversity analysis, the haplotype diversity (H_d_), nucleotide diversity (π), number of segregating sites (S), mutations parameter based on segregating sites (Sθ) and the mean number of pairwise differences (k) values within the aligned consensus sequences of *P. vivax dhfr* locus were calculated using the DnaSP 5.10 software package (Librado and Rozas, 2009). For the analysis of genetic differences, the pairwise fixation index (*F*_ST_) values between the aligned *P. vivax dhfr* consensus sequences were calculated using Arlequin program v. 3.5.2.2 (Loftus et al., 1994).

For phylogenetic analysis, the aligned *P. vivax dhfr* consensus sequences were imported into the FaBox 1.5 online tool to collapse all sequences showing 100% base pair similarity after corrections into a single haplotype (unique sequences generated from millions of MiSeq reads). A split tree of *dhfr* haplotypes was constructed based on the HKY85 genetic model using the neighbour-joining method employed in SplitTrees4 software v4.10 (Huson & Bryant, 2006). The appropriate model of nucleotide substitutions for neighbour-joining analysis was selected by using the jModeltest 12.2.0 program (Posada, 2008).

## 3. Results

### 3.1. Allele frequencies of pyrimethamine resistance-associated SNPs

A 468 bp fragment of *P. vivax dhfr* locus was successfully amplified from 141 individual isolates selected from our previous study (Wahab et al., 2020). The relative allele frequencies of five resistance-associated SNPs [F57L/I (TT<u>C</u>-TT<u>A</u>/<u>A</u>T<u>A</u>), S58R (AG<u>C</u>-AG<u>A</u>), T61M (A<u>C</u>G-A<u>T</u>G), S117N/T (A<u>G</u>C-A<u>A</u>C/A<u>C</u>C) and I173L/F (A<u>T</u>T-<u>C</u>TT/<u>T</u>TT)] were determined by metabarcoded deep amplicon sequencing technology. Ninty-nine *P. vivax* isolates were 100% susceptible based on the *dhfr* locus allele frequencies [F57 (TT<u>C</u>), S58 (AG<u>C</u>), T61 (A<u>C</u>G), S117 (A<u>G</u>C) and I173 (A<u>T</u>T)] (Fig. 1). Pyrimethamine susceptible alleles and resistance-associated mutations were present in 42 *P. vivax* isolates (Fig. 1, Table 1). Based on the analysis of each isolate separately, 11 isolates carried between 62.44 and 98.95% susceptible alleles and 1.05 and 37.56% carried resistance mutations [Table 1, Isolate ID (R-1-R-11)], and 31 isolates carried between 51.98 and 100% resistant-associated mutations and 0 and 48.02% susceptible alleles [Table 1, Isolate ID (R-12-R-42)]. Based on the analysis of each resistance SNP separately, 117N (A<u>A</u>C) single mutation was identified in 23 isolates at frequencies between 0.04 and 100%. Double mutations of 58R/117N (AG<u>A</u>/A<u>A</u>C) were detected in 33 isolates at frequencies between 0.06 and 100%. The 173L (<u>C</u>TT) SNP was identified in 4 isolates at frequencies between 0.99 and 100% (Fig. 1, Table 1). The single pyrimethamine resistance-associated mutation at 58R (AG<u>A</u>) was identified in 3 isolates at low frequencies between 0.06 and 0.74% (Fig 1, Table 1). Triple mutations of 57L/58R/117N (TT<u>A/</u>AG<u>A</u>/A<u>A</u>C), 58R/61M/117N (AG<u>A</u>/A<u>T</u>G/A<u>A</u>C) and 58R/117N/173L (AG<u>A</u>/A<u>A</u>C/<u>C</u>TT) were present in 4, 1 and 1 isolates respectively, with very low frequencies of 0.06% (Fig 1, Table 1).

**Table 1:** Relative allele frequencies of the *dhfr* pyrimethamine resistance-associated mutations in 42 *P. vivax* isolates showing resistance reads, from the Punjab province of he relative allele frequency of resistance versus susceptible was based on the alleles identification using Illumina MiSeq deep amplicon sequencing technology.

| Isolates ID | Total no. of Illumina Miseq reads | Total no of susceptible reads | Total no of resistant reads | Susceptible alleles | 58R/117N resistant alleles | 58R resistant alleles | 117N resistant alleles | 173L resistant alleles | 57L//S58R/117N resistant alleles | 58R/61M/117N resistant alleles | 58R/117N/173L resistant alleles | Region |
| --- | --- | --- | --- | --- | --- | --- | --- | --- | --- | --- | --- | --- |
| R-1-34 | 3528 | 3095 | 433 | 98.95 | 0.06 |  |  | 0.99 |  |  |  | Lahore |
| R-2-32 | 1402 | 1179 | 223 | 98.09 |  |  |  | 1.91 |  |  |  | Lahore |
| R-3-139 | 2092 | 1767 | 325 | 98.60 | 1.40 |  |  |  |  |  |  | Lahore |
| R-4-141 | 3148 | 2384 | 764 | 95.45 | 4.55 |  |  |  |  |  |  | Lahore |
| R-5-10 | 1880 | 1548 | 332 | 94.48 |  |  | 5.52 |  |  |  |  | D.G. Khan |
| R-6-45 | 9265 | 7383 | 1882 | 91.41 |  |  | 8.59 |  |  |  |  | Larkana |
| R-7-92 | 3084 | 2423 | 661 | 75.00 | 25.00 |  |  |  |  |  |  | Faisalabad |
| R-8-84 | 3542 | 2406 | 1136 | 67.93 | 31.17 |  | 0.90 |  |  |  |  | D.G.Khan |
| R-9-27 | 5615 | 4576 | 1039 | 66.09 | 33.91 |  |  |  |  |  |  | D.G.Khan |
| R-10-104 | 8296 | 6766 | 1530 | 63.27 | 34.39 | 0.06 | 2.27 |  |  |  |  | Lahore |
| R-11-105 | 9325 | 7120 | 2205 | 62.44 | 36.35 | 0.74 | 0.47 |  |  |  |  | Lahore |
| R-12-26 | 6657 | 3025 | 3632 | 45.44 | 51.98 | 0.12 | 2.46 |  |  |  |  | Lahore |
| R-13-128 | 8729 | 2508 | 6221 | 37.97 | 61.30 |  | 0.73 |  |  |  |  | Lahore |
| R-14-102 | 2192 | 533 | 1659 | 30.64 | 69.31 |  | 0.04 |  |  |  |  | Sheikhupura |
| R-15-109 | 7647 | 1211 | 6436 | 14.29 | 85.71 |  |  |  |  |  |  | D.G.Khan |
| R-16-55 | 5128 | 792 | 4336 | 7.59 |  |  | 92.41 |  |  |  |  | Lahore |
| R-17-33 | 6993 | 566 | 6427 | 5.39 | 94.10 |  | 0.52 |  |  |  |  | D.G.Khan |
| R-18-39 | 2656 | 234 | 2422 | 2.34 | 93.89 |  |  | 3.78 |  |  |  | Lahore |
| R-19-126 | 3362 |  | 3362 |  | 99.07 |  | 0.10 |  |  |  |  | Rahem Yar Khan |
| R-20-83 | 3608 |  | 3608 |  | 99.70 |  | 0.03 |  |  |  |  | Lahore |
| R-21-14 | 9040 |  | 9040 | 0.2 |  |  | 100 |  |  |  |  | Layyah |
| R-22-121 | 7658 |  | 7658 |  | 99.80 |  | 0.03 |  |  |  |  | D.G.Khan |
| R-23-82 | 4625 |  | 4625 |  | 99.78 |  | 0.01 |  | 0.06 |  |  | Lahore |
| R-24-78 | 3847 |  | 3847 |  | 99.89 |  |  |  |  |  |  | Lahore |
| R-25-118 | 6697 |  | 6697 |  | 2.12 |  | 97.82 |  |  |  |  | Lahore |
| R-26-63 | 5657 |  | 5657 |  | 99.93 |  |  |  | 0.02 |  |  | Lahore |
| R-27-132 | 4900 |  | 4900 |  | 99.93 |  | 0.03 |  |  |  |  | Lahore |
| R-28-93 | 2889 |  | 2889 |  | 99.95 |  | 0.03 |  |  |  |  | Lahore |
| R-29-97 | 3212 |  | 3212 |  |  |  | 100 |  |  |  |  | Sheikhupura |
| R-30-130 | 9325 |  | 9325 |  | 99.98 |  | 0.02 |  |  |  |  | Lahore |
| R-31-101 | 8495 |  | 8495 |  | 99.97 |  |  |  | 0.01 | 0.01 | 0.01 | Peshawar |
| R-32-88 | 4333 |  | 4333 |  | 99.98 |  |  |  | 0.02 |  |  | Lahore |
| R-33-17 | 7447 |  | 7447 |  | 99.99 |  | 0.01 |  |  |  |  | Lahore |
| R-34-142 | 3219 |  | 3219 |  | 100 |  |  |  |  |  |  | Lahore |
| R-35-112 | 7342 |  | 7342 |  | 100 |  |  |  |  |  |  | Larkana |
| R-36-7 | 2412 |  | 2412 |  | 100 |  |  |  |  |  |  | Gujranwala |
| R-37-37 | 2115 |  | 2115 |  | 100 |  |  |  |  |  |  | Lahore |
| R-38-51 | 3110 |  | 3110 |  | 100 |  |  |  |  |  |  | Lahore |
| R-39-64 | 1869 |  | 1869 |  | 100 |  |  |  |  |  |  | Lahore |
| R-40-53 | 4836 |  | 4836 |  |  |  | 100 |  |  |  |  | Lahore |
| R-41-131 | 3278 |  | 3278 |  |  |  | 100 |  |  |  |  | Lahore |
| R-42-28 | 2831 |  | 2831 |  |  |  |  | 100 |  |  |  | Sheikhupura |

**Fig. 1.**
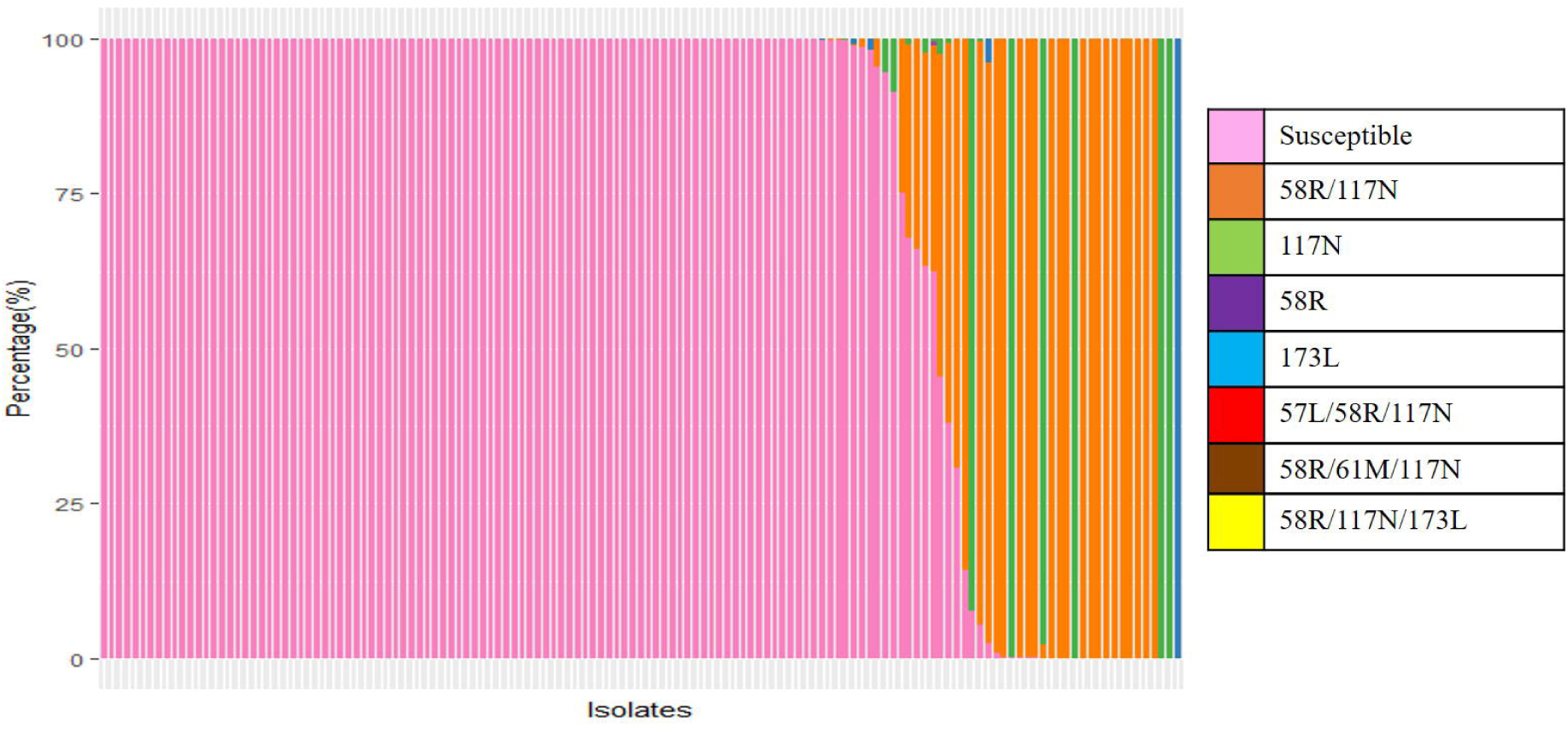
Relative allele frequencies of the pyrimethamine resistance-associated mutations in 141 *P*.*vivax* isolates from the Punjab province of Pakistan. The frequency of resistance and susceptible alleles was based on their identification using Illumina MiSeq deep amplicon sequencing technology. Resistant and susceptible alleles are shown in different colours.

### 3.2. Phylogeny of pyrimethamine resistance-associated SNPs

A total of 178 unique *dhfr* haplotypes were generated among 42 *P. vivax* isolates carried pyrimethamine susceptible and resistance-associated mutations (Supplementary Data S1). Seventy-four out of 178 haplotypes [F57 (TT<u>C</u>), S58 (AG<u>C</u>), T61 (A<u>C</u>G), S117 (A<u>G</u>C) and I173 (A<u>T</u>T)] were susceptible, and 104 out of 178 haplotypes [58R (AG<u>A</u>), 117N (A<u>A</u>C) and 173N (A<u>A</u>C)] were resistant. Out of those 104 haplotypes, 54 encoded 58R/117N (AG<u>A</u>/A<u>A</u>C) double resistance mutations, 40 encoded 117N (A<u>A</u>C), 5 encoded 58R (AG<u>A</u>) and 2 encoded 173L (<u>C</u>TT) resistance mutations. Three haplotypes encoded the triple resistance mutations of 57L/58R/117N (TT<u>A/</u>AG<u>A</u>/A<u>A</u>C), 58R/61M/117N (AG<u>A</u>/A<u>T</u>G/A<u>A</u>C) and 58R/117N/173L (AG<u>A</u>/A<u>A</u>C/<u>C</u>TT) (Supplementary Data S1).

A split tree was created to examine the phylogenetic relationship between 104 unique *dhfr* resistance haplotypes identified among 42 *P. vivax* isolates (Fig. 2). The analysis of the 58R/117N (AG<u>A</u>/A<u>A</u>C) resistance mutants showed that 54 haplotypes were located in a single lineage (Fig. 2). Three individual haplotypes of the triple mutants [57L/58R/117N (TT<u>A</u>/AG<u>A</u>/A<u>A</u>C), 58R/61M/117N (AG<u>A</u>/A<u>T</u>G/A<u>A</u>C), 58R/117N/173L (AG<u>A</u>/A<u>A</u>C/<u>C</u>TT)] shared the lineage of 58R/117N (AG<u>A</u>/AAC) haplotypes (Fig 2). Analysis of the 117N (A<u>A</u>C) resistance mutant revealed that 40 haplotypes were located in 2 separate lineages (Fig. 2). Five haplotypes of 58R (AG<u>A</u>) and 2 haplotypes of 173L (<u>C</u>TT) mutant were found at low frequencies in two separate lineages (Fig. 2).

**Fig. 2.**
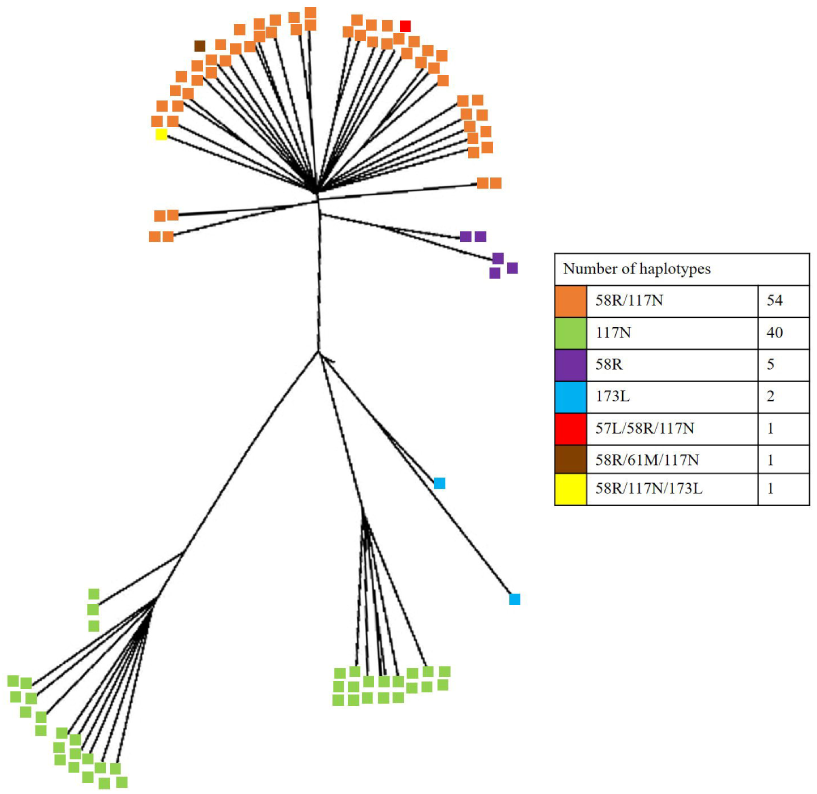
A split tree was generated from 104 haplotypes identified in *P. vivax dhfr* sequence data (Supplementary Data S1). The haplotypes were aligned on the MUSCLE tool of the Geneious v9.0.1 and the tree was constructed with the UPGMA method in the HKY85 model of substitution in the SplitsTrees4 software. The appropriate model of nucleotide substitutions was selected by using the jModeltest 13.1.0 program. The circles in the tree represent different mutations in *dhfr* locus containing different colours.

### 3.3. Genetic structure of dhfr locus

The genetic structure of the *dhfr* locus was assessed individually from 42 *P. vivax* isolates. The data show a high genetic diversity at both haplotype and nucleotide levels (Table 2), with the values of haplotype diversity (H_d_) ranging from 0.111 to 0.923 and nucleotide diversity (π) from 0.00021 to 0.00598 within individual isolate. The overall values between isolates were 0.769 and 0.00415 for haplotype and nucleotide diversity, respectively. The mean fixation index (F_ST_) values of the *dhfr* locus were 0.62, indicating low to moderate genetic differentiation within 42 *P. vivax* isolates. The F_ST_ values also indicated a low to moderate genetic differentiation, ranging from 0.01 to 0.98 between individual isolate (Fig. 3).

**Table 2:** Summary of the genetic diversity data for the *dhps* locus of 42 *P. vivax* isolates.

| Isolates ID | (H<sub>d</sub>) | (S) | ($\Pi$) | ($\Theta_s$) | (k) |
| --- | --- | --- | --- | --- | --- |
| R-1-34 | 0.111 | 1 | 0.00025 | 0.00066 | 0.121 |
| R-2-32 | 0.294 | 1 | 0.00066 | 0.00065 | 0.294 |
| R-3-139 | 0.659 | 3 | 0.00258 | 0.00213 | 1.143 |
| R-4-141 | 0.682 | 4 | 0.00428 | 0.00297 | 1.909 |
| R-5-10 | 0.173 | 1 | 0.00042 | 0.00067 | 0.173 |
| R-6-45 | 0.390 | 5 | 0.00110 | 0.00282 | 0.468 |
| R-7-92 | 0.538 | 2 | 0.00242 | 0.00141 | 1.077 |
| R-8-84 | 0.600 | 3 | 0.00320 | 0.00216 | 1.371 |
| R-9-27 | 0.500 | 2 | 0.00224 | 0.00016 | 1.000 |
| R-10-104 | 0.439 | 4 | 0.00197 | 0.00202 | 0.767 |
| R-11-105 | 0.753 | 5 | 0.00287 | 0.00288 | 1.150 |
| R-12-26 | 0.788 | 4 | 0.00380 | 0.00297 | 1.697 |
| R-13-128 | 0.800 | 5 | 0.00456 | 0.00338 | 2.033 |
| R-14-102 | 0.857 | 6 | 0.00440 | 0.00389 | 1.886 |
| R-15-109 | 0.545 | 2 | 0.00255 | 0.00155 | 1.091 |
| R-16-55 | 0.882 | 4 | 0.00369 | 0.00317 | 1.644 |
| R-17-33 | 0.818 | 5 | 0.00371 | 0.00371 | 2.318 |
| R-18-39 | 0.729 | 6 | 0.00598 | 0.00388 | 2.571 |
| R-19-126 | 0.383 | 3 | 0.00141 | 0.00183 | 0.625 |
| R-20-83 | 0.228 | 2 | 0.00076 | 0.00133 | 0.338 |
| R-21-14 | 0.471 | 1 | 0.00108 | 0.00167 | 0.471 |
| R-22-121 | 0.775 | 17 | 0.00388 | 0.00188 | 1.664 |
| R-23-82 | 0.288 | 4 | 0.00096 | 0.00224 | 0.425 |
| R-24-78 | 0.200 | 2 | 0.00090 | 0.00159 | 0.400 |
| R-25-118 | 0.754 | 5 | 0.00378 | 0.00312 | 1.623 |
| R-26-63 | 0.145 | 3 | 0.00050 | 0.00175 | 0.222 |
| R-27-132 | 0.251 | 4 | 0.00088 | 0.00229 | 0.386 |
| R-28- 93 | 0.923 | 11 | 0.00496 | 0.00804 | 2.132 |
| R-29-97 | 0.451 | 10 | 0.00131 | 0.00521 | 0.546 |
| R-30-130 | 0.243 | 6 | 0.00073 | 0.00335 | 0.308 |
| R-31-101 | 0.539 | 16 | 0.00162 | 0.00754 | 0.681 |
| R-32-88 | 0.143 | 1 | 0.00033 | 0.00073 | 0.143 |
| R-33-17 | 0.541 | 3 | 0.00141 | 0.00196 | 0.637 |
| R-34-142 |  |  | N/A |  |  |
| R-35-112 | 0.868 | 7 | 0.00278 | 0.00494 | 1.242 |
| R-36-7 |  |  | N/A |  |  |
| R-37-37 |  |  | N/A |  |  |
| R-38-51 | 0.195 | 1 | 0.00021 | 0.00063 | 0.095 |
| R-39-64 |  |  | N/A |  |  |
| R-40-53 | 0.121 | 1 | 0.00065 | 0.00025 | 0.098 |
| R-41-131 | 0.172 | 5 | 0.00050 | 0.00258 | 0.222 |
| R-42-28 |  |  | N/A |  |  |
| Total | 0.769 | 38 | 0.00415 | 0.01385 | 1.523 |

Haplotype diversity (H<sub>e</sub>); the number of segregating sites (S); nucleotide diversity ( $\pi$ ); the mean number of pairwise differences (k); the mutation parameter based on an infinite site equilibrium model, and the mutations parameter based on segregating sites (S<sub>0</sub>).

**Fig. 3.**
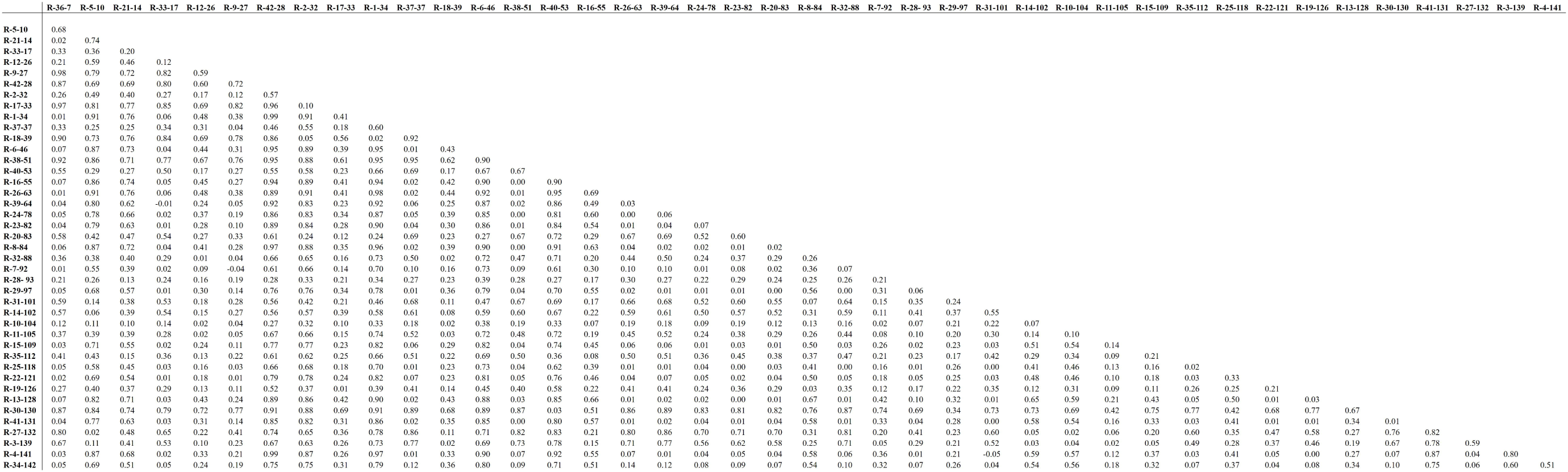
Fixation index (F_ST_) values based on genotyping of 42 *P. vivax* isolates using *dhfr* sequence data.

## 4. Discussion

Several studies have explored the origins of pyrimethamine resistance-associated mutations in *P. falciparum* in different geographical regions. *P. falciparum dhfr* quartet mutants have been identified with a single genetic origin in Southeast Asian countries (Mita et al., 2009). In contrast, *P. falciparum dhfr* triple and double mutants have been identified with multiple origins in Southeast Asia, South America and African countries (Lumb et al., 2009; Mita, 2010). Understanding the nature of adaptive changes associated with the origin of pyrimethamine resistance in *P. vivax* is poor (Hawkins et al., 2008b).

In the present study, 54 diverse haplotypes of the 58R/117N double mutants were present on a single lineage, suggesting that there is a single origin of this mutation in *P. vivax* isolates examined. The triple mutants (57L/58R/117N, 58R/61M/117N and 58R/117N/173L) shared this lineage, suggesting that these mutations have the same origins. Five haplotypes of 58R and 2 haplotypes of 173L mutants were found at low frequencies in two separate lineages, indicating single origin. Forty diverse haplotypes of the 117N mutant were present on two separate lineages suggesting multiple origins of this mutation. These results differ from those of a previous study of the evolutionary origin of *P. vivax dhfr* resistance-conferring mutations (Hawkins et al., 2008b), which demonstrated that 58R/117N double mutants, 58R/61M/117T triple mutants and 57L/61M/117T/173F, 57I/58R/61M/117T, 57L/58R/61M/117T quadruple mutants had multiple origins in Thailand, Indonesia, Papua New Guinea and Vanuatu. A level of genetic diversity in *P. vivax*, may confer genetic adaptability (Alam et al., 2007; Hong et al., 2016), enabling the origin of pyrimethamine resistance mutations. In the present study, we have identified a high level of allelic polymorphism in *P. vivax* isolates, consistent with the high level of genetic diversity expected for this parasite. Conversely, a high mutation rate (2.5 × 10^−9^) was shown in *P. falciparum* in an experiment measuring mutations associated with the origin of pyrimethamine resistance mutations (Paget-McNicol and Saul, 2001). The effective parasite load may also influence the origin of pyrimethamine resistance mutations in *P. vivax* (Hastings et al., 2004).

The genetic differentiation at the *dhfr* locus amongst the *P. vivax* isolates for the current study was consistent with human migration between the cities of the Punjab province of Pakistan playing a role in the spread of pyrimethamine resistance mutations. The spread of resistance mutations may be influenced by the impact of the antimalarial drug on the gametocytes stage of *Plasmodium*. It has been demonstrated that pyrimethamine resistance can increase the number of gametocytes carried by the patient, thereby increasing transmission intensity of resistant parasites during a mosquito blood meal (Petersen et al., 2011).

The present study describes the allele frequencies of pyrimethamine resistance mutations in the *dhfr* locus of *P. vivax* isolates from the Punjab Province of Pakistan. Our findings are similar to previous studies from Pakistan, where a 117N single mutant and 58R/117N double mutants were shown to be highly prevalent, whilst the 57L, 58R and 61M mutants were only detected at low frequencies and or, in combination with the 117N mutant (Khattak et al., 2013; Raza et al., 2013; Shaukat et al., 2019; Zakeri et al., 2011). Previous studies have consistently shown that the 117N single mutant and 58R/117N/T double mutants were present at high frequencies in different geographical regions, while the 57L, 58R, 61M, and 173L single mutants and 57L/58R/117N, 58R/61M/117N and 58R/117N/173L triple mutants were present at relatively low frequencies (Auliff et al., 2006; Brega et al., 2004; de Pecoulas et al., 1998; Hastings et al., 2005; Imwong et al., 2003; Kaur et al., 2006; Kuesap et al., 2011; Lu et al., 2012; Mint Lekweiry et al., 2012; Ranjitkar et al., 2011; Schunk et al., 2006). Mutations in the *P. vivax dhfr* locus may impart a fitness cost, whereby the selective advantage acquired by becoming drug-resistant is balanced by the biological cost arising from the altered function of the mutated protein (Petersen et al., 2011). The single origins of the common mutants and single origins of the rare mutants shown in the present study may reflect differences in fitness costs between these mutations (Petersen et al., 2011).

In conclusion, we investigated evidence for multiple and single origins of different SNPs in the *dhfr* locus of *P. vivax* associated with pyrimethamine resistance. The results show high allele frequencies associated with 58R/117N and 117N resistance mutations and relatively low frequencies of other mutations. Understanding the origin of resistance mutations is needed to develop strategies for prolonging the effectiveness of pyrimethamine drug treatment. From these findings, better surveillance methods can be established to monitor the dispersion of the pyrimethamine resistance.

## Acknowledgement

Work at the University of Central Punjab, Pakistan and Kohat University of Science and Technology Pakistan uses facilities funded by the Higher Education Commission of Pakistan.

